# LKB1 Is Physiologically Required for Sleep from Drosophila to the Mouse

**DOI:** 10.1101/2021.12.13.472078

**Authors:** Ziyi Liu, Lifen Jiang, Chaoyi Li, Chengang Li, Jingqun Yang, Dan Wang, Renbo Mao, Yi Rao

## Abstract

LKB1 is known as a master kinase for 14 kinases related to the adenosine monophosphate (AMP)-activated protein kinase (AMPK). Two of them (SIK3 and AMPKα) have previously been implicated in sleep regulation. We generated loss-of-function (LOF) mutants for Lkb1 in both Drosophila and mice. Sleep was reduced in Lkb1-mutant flies and in flies with neuronal deletion of Lkb1. Sleep was reduced in mice after virally mediated reduction of Lkb1 in the brain. Electroencephalography (EEG) analysis showed that non-rapid eye movement (NREM) sleep and sleep need were both reduced in Lkb1-mutant mice. These results indicate that LKB1 plays a physiological role in sleep regulation conserved from flies to mice.

## INTRODUCTION

Human Peutz-Jeghers syndrome (PJS) (Jeghers et al. 1949; Peutz 1921) is an autosomal dominant disorder with gastrointestinal (GI) polyps and increased risk of multiple tissues (Tomlinson and Houlston 1997; Westerman et al. 1999). The gene mutated in, and responsible for, PJS encodes the liver kinase B1 (LKB1, also known as STK11) (Hemminki et al. 1998; Hemminki et al. 1997; Jenne et al. 1998). Lkb1 is thus a tumor suppressor gene, mutated in multiple cancers, especially the GI (Bardeesy et al. 2002; Hearle et al. 2006; Jishage et al. 2002; Mehenni et al. 1998; Miyoshi et al. 2002) and lung adenocarcinoma (Carretero et al. 2004; Gill et al. 2011; Ji et al. 2007; Matsumoto et al. 2007; Sanchez-Cespedes et al. 2002; Skoulidis et al. 2015), cervical cancer (Wingo et al. 2009), ovarian cancer (Tanwar et al. 2014), breast cancer (Hearle et al. 2006; Sengupta et al. 2017; Shen et al. 2002), pancreatic cancer (Morton et al. 2010) and melanoma (Guldberg et al. 1999; Rowan et al. 1999).

LKB1 phosphorylates threonine 172 (T172) of the α subunit of adenosine monophosphate (AMP)-activated protein kinase (AMPKα) (Hawley et al. 2003; Hong et al. 2003; Lizcano et al. 2004; Sakamoto et al. 2005; Shaw et al. 2004; Shaw et al. 2005; Sutherland et al. 2003; Woods et al. 2003), and positively regulates the activity of AMPK.

AMPK is a well-known kinase (Beg et al. 1973; Carling et al. 1989; Carling et al. 1987; Carlson and Kim 1973; Ferrer et al. 1985; Ingebritsen et al. 1978; Munday et al. 1988; Yeh and Kim 1980) with important physiological and pathological roles (Hardie 2014; Hardie et al. 2016; Herzig and Shaw 2018; Lopez and Dieguez 2014). The α, β and γ subunits of AMPK form a heterotrimeric complex (Davies et al. 1994; Michell et al. 1996; Mitchelhill et al. 1994). The catalytic α subunit is regulated by phosphorylation at T172 of AMPKα2 or T183 of AMPKα1 (Hawley et al. 1996).

There are 12 additional mammalian AMPK-related kinases (ARKs), similar to the α subunit of AMPK, all regulated at the site equivalent to AMPK-T172 (Lizcano et al. 2004). LKB1 and its associated proteins STE20-related adaptor (STRAD) and mouse protein 25 (MO25) have been reported to phosphorylate all 14 ARKs (Lizcano et al. 2004), making LKB1 a master kinase for ARKs (Alessi et al. 2006; Lizcano et al. 2004; Shackelford and Shaw 2009).

Some ARKs have been reported to regulate sleep. In mice, inhibitors of AMPK were found to decrease sleep, whereas activators of AMPK were found to increase sleep (Chikahisa et al. 2009). In flies, knockdown of AMPKβ in neurons decreased the total amount of sleep and resulted in fragmented sleep (Nagy et al. 2018). Knockdown of AMPKα in a specific pair of neurons suppressed sleep (Yurgel et al. 2019).

Studies in mice have shown that sleep was increased in gain-of-function (GOF) mutations in the salt inducible kinase (SIK) 3 (Funato et al. 2016), and sleep need was reduced in GOF mutants of SIK 1, 2 and 3 (Funato et al. 2016; Honda et al. 2018; Park et al. 2020). Sleep was also decreased when SIK3 was downregulated in flies (Funato et al. 2016).

Here we investigated the functional role of LKB1 in regulating sleep in flies and mice.

## RESULTS

### Sleep Phenotypes of Lkb1 Mutant Drosophila

Null mutants for Lkb1 are lethal in Drosophila (Martin and St Johnston 2003). We had generated a Lkb1 knockout (“lkb1^T1^”) line (Figure S1A, S2B) and found that lkb1^T1/T1^ mutation was lethal in the pupa stage. The level of Lkb1 mRNA was reduced in the heterozygous lkb1^T1/+^ flies (Figure S1B). We then tested whether the heterozygous lkb1^T1^ had any phenotype in sleep using flies kept in 12 hours (h) light/ 12 h dark (LD) cycles (Figure S1C). While lkb1^T1/+^ flies were not significantly different from the wild type (wt) flies in sleep bout numbers (Figure S1E), or daytime sleep duration (Figure S1D), daytime sleep bout duration (Figure S1F), lkb1^T1/+^ flies showed significantly lower nighttime sleep duration (Figure S1D), nighttime sleep bout duration (Figure S1F) and longer latency to sleep (Figure S1G). Thus, there was dosage-sensitive physiological requirement of Lkb1 in nighttime sleep.

We tried to, and succeeded in, generating lkb1^T2^, a hypomorphic mutation for Lkb1 in flies (Figure 1A, S2A and S2B). Lkb1 mRNA was significantly reduced in lkb1^T2/+^ and lkb1^T2/T2^ flies (Figure 1B).

**Figure 1.**
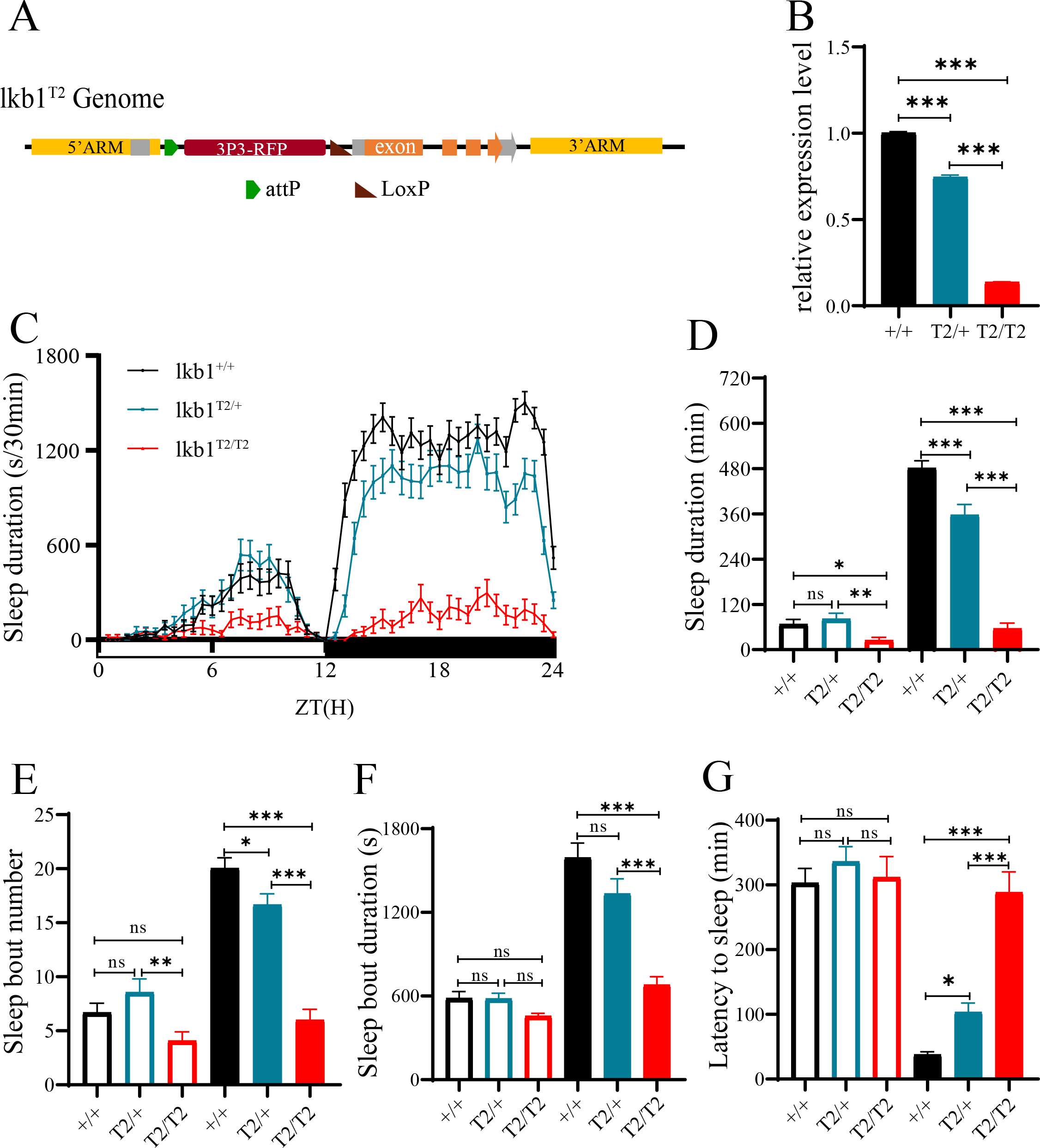
**Sleep Phenotypes of Lkb1 Knock-Down Mutants.** (A) A diagram illustrating the Lkb1 insertion mutant lkb1^T2^. (B) Relative Lkb1 mRNA levels in lkb1^T2/T2^ (red), lkb1^T2/+^ (blue) and wt (black) flies. (C) Sleep profiles of lkb1^T2/T2^ (red, n=42), lkb1^T2/+^ (blue, n=44), and wt (black, n=44) flies in a 12 h light/12 h dark (LD) cycle. (D-G) Statistical analysis of sleep duration, sleep bout number, sleep bout duration and latency to sleep in lkb1^T2/T2^ (red, n=42), lkb1^T2/+^ (blue, n=44) and wt (black, n=44) flies. Open bars denote daytime, filled bars denote nighttime. (D) Sleep duration. Daytime and nighttime sleep durations of lkb1^T2/T2^ mutants was significantly less than those in lkb1^T2/+^ and wt flies. (E) Sleep bout number. Daytime sleep bout number of lkb1^T2/T2^ mutants was less than that of lkb1^T2/+^ flies. Nighttime sleep bout number of lkb1^T2/T2^ was significantly less than those of lkb1^T2/+^ and wt flies. (F) Sleep bout duration. Nighttime sleep bout duration of lkb1^T2/T2^ mutants was significantly less than those of lkb1^T2/+^ and wt flies. (G) Latency to sleep. Latency to sleep after light-off of lkb1^T2/T2^ mutants was significantly prolonged than lkb1^T2/+^ and wt flies. Ordinary One-way ANOVA was used. n.s. denotes p>0.05, *p<0.05, **p<0.01, ***p<0.001. Error bars represent standard error of the mean (SEM).

During the day, sleep duration was significantly less in lkb1^T2/T2^ flies than in lkb1^T2/+^ and wt flies (Figure 1C, 1D), sleep bout number was significantly less in lkb1^T2/T2^ flies than in lkb1^T2/+^ and wt flies (Figure 1E), while sleep bout duration and latency showed no difference in three groups (Figure 1F, 1G).

During the night, not only lkb1^T2/T2^ flies showed highly reduced sleep duration (Figure 1C, 1D), highly reduced sleep bout number (Figure 1E), highly reduced sleep bout duration (Figure 1F) and highly increased latency (Figure 1G) than the wt flies, but the lkb1^T2/+^ flies were also significantly different from the wt flies in all these parameters (Figure 1C to 1G), confirming a dosage sensitive requirement for Lkb1. Results of sleep analysis of lkb1^T1/+^, lkb1^T2/+^ and lkb1^T2/T2^ mutant flies are all consistent with the possibility that Lkb1 plays a physiological role in promoting sleep.

### Rescue of Sleep Phenotypes by Lkb1 in Flies

We inserted the sequence of the yeast transcription factor Gal4 into the lkb1^T2^ mutant flies, flanking the lkb1 promoter, and obtained lkb1^T2^-Gal4 flies (Figure 2A). We also generated UAS-Lkb1 flies in which the Lkb1 coding sequence (CDS) was expressed under the control of the upstream activation sequence (UAS). Because Gal4 protein binds to the UAS (Brand and Perrimon 1993), the expression of Lkb1 in flies resulting from the crosses between lkb1^T2^-Gal4 flies and UAS-Lkb1 flies would be under the control of the endogenous Lkb1 promoter. Indeed, expression of Lkb1 mRNA was restored when lkb1^T2^-Gal4 and UAS-Lkb1 were present in the same flies (Figure 2B), whereas Lkb1 mRNA was less in UAS-Lkb1;lkb1^T2/T2^ mutant flies, and lkb1^T2^-Gal4/ lkb1^T2^-Gal4 flies than that in the wt. UAS-Lkb1 alone could not restore Lkb1 mRNA expression level to that in wt flies (Figure 2B).

**Figure 2.**
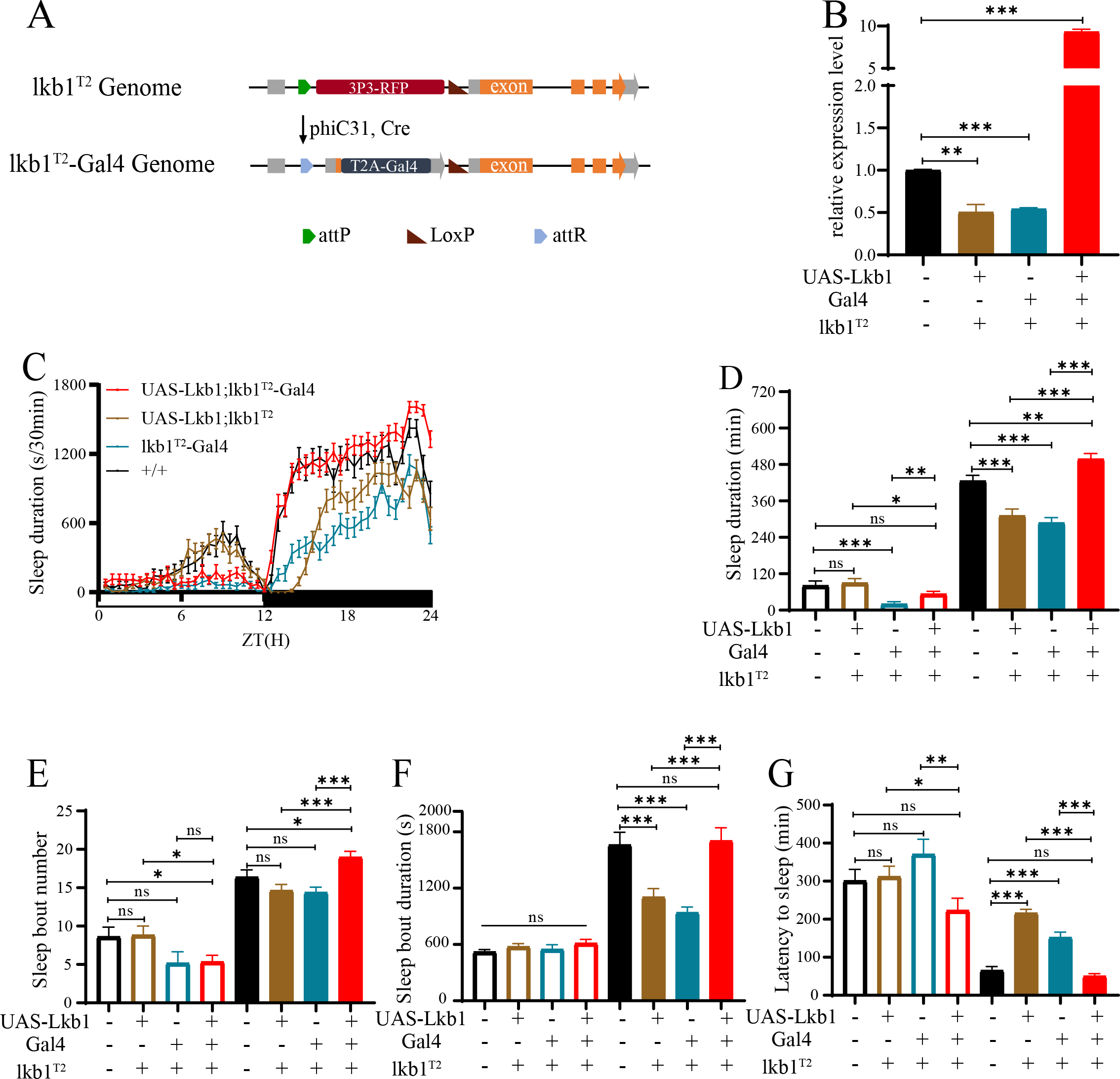
**Rescue of Sleep Loss in lkb1^T2/T2^ by Lkb1.** (A) A diagram of lkb1^T2^-Gal4: a cDNA for the yeast Gal4 gene inserted in the lkb1^T2^ knockdown mutants. (B) Relative Lkb1 mRNA levels in lkb1^T2/T2^-Gal4 (blue), UAS-Lkb1; lkb1^T2/T2^-Gal4 (red), UAS-Lkb1; lkb1^T2/T2^ (yellow) and wt (black) flies. (C-G) In lkb1^T2/T2^-Gal4 homozygous flies, UAS-Lkb1 cDNA driven by Gal4 to rescue sleep phenotypes of lkb1 knockdown mutants. (C) Sleep profiles of UAS-Lkb1; lkb1^T2/T2^-Gal4 (red, n=45), UAS-Lkb1; lkb1^T2/T2^ (yellow, n=47), lkb1^T2/T2^-Gal4 (blue, n=46) and wt (black, n=36) flies. (D-G) Statistical analysis of sleep duration, sleep bout number, sleep bout duration and latency to sleep in UAS-Lkb1; lkb1^T2/T2^-Gal4 (red, n=45), UAS-Lkb1; lkb1^T2/T2^ (yellow, n=47), lkb1^T2/T2^-Gal4 (blue, n=46) and wt (black, n=36) flies. Open bars denote daytime, filled bars nighttime. (D) Sleep duration. Nighttime sleep duration of UAS-Lkb1; lkb1^T2/T2^-Gal4 was similar to that of wt mutants, both significantly higher than UAS-Lkb1; lkb1^T2/T2^ and lkb1^T2/T2^-Gal4 flies. (E) Sleep bout number. Nighttime sleep bout number of UAS-Lkb1; lkb1^T2/T2^-Gal4 was similar to the wt but significantly higher than UAS-Lkb1; lkb1^T2/T2^ and lkb1^T2/T2^-Gal4 flies. (F) Sleep bout duration. Nighttime sleep bout duration of UAS-Lkb1; lkb1^T2/T2^-Gal4 was similar to the wt but significantly higher than UAS-Lkb1; lkb1^T2/T2^ and lkb1^T2/T2^-Gal4 flies. (G) Latency to sleep. Latency after light-off of UAS-Lkb1; lkb1^T2/T2^-Gal4 was similar to the wt but significantly shorter than UAS-Lkb1; lkb1^T2/T2^ and lkb1^T2/T2^-Gal4 flies. Unpaired t test was used. n.s. denotes p>0.05, * p<0.05, ** p<0.01, ***p<0.001. Error bars represent SEM.

Both daytime and nighttime sleep durations were less in lkb1^T2^-Gal4/ lkb1^T2^-Gal4 flies than those in wt flies (Figure 2C). Introduction of UAS-Lkb1 in lkb1^T2/T2^ flies or lkb1^T2^-Gal4 alone could not restore sleep. When both lkb1^T2^-Gal4 and UAS-Lkb1 were present, nighttime sleep durations were restored (Figure 2D). Nighttime sleep bout number, nighttime sleep bout duration and nighttime latency were restored when both lkb1^T2^-Gal4 and UAS-Lkb1 were present, but not when lkb1^T2^-Gal4 or UAS-Lkb1 alone was present (Figure 2E, 2F and 2G).

These results support that the sleep phenotypes of lkb1^T2/T2^ were resulted from the reduction of Lkb1 mRNA expression in these flies.

### Sleep Phenotypes of Flies Carrying Neuronal Deletion of the Lkb1 Gene

To determine whether Lkb1 functions in neurons, we used the CRISPR-Cas9 system to delete Lkb1 from neurons specifically (Figure S3). A pan-neuronal Gal4 driver (57C10-Gal4) was used to control the expression of small guide RNA (sgRNA) targetting Lkb1 in neurons. Sleep was not significantly different when either 57C10-Gal4 or UAS-Lkb1 sgRNA alone was introduced into flies (Figure 3), but when both 57C10-Gal4 and UAS-Lkb1 sgRNA were present in flies, nighttime sleep duration (Figure 3B) and nighttime sleep bout duration (Figure 3D) were significantly reduced and nighttime sleep latency significantly lengthened (Figure 3E).

**Figure 3.**
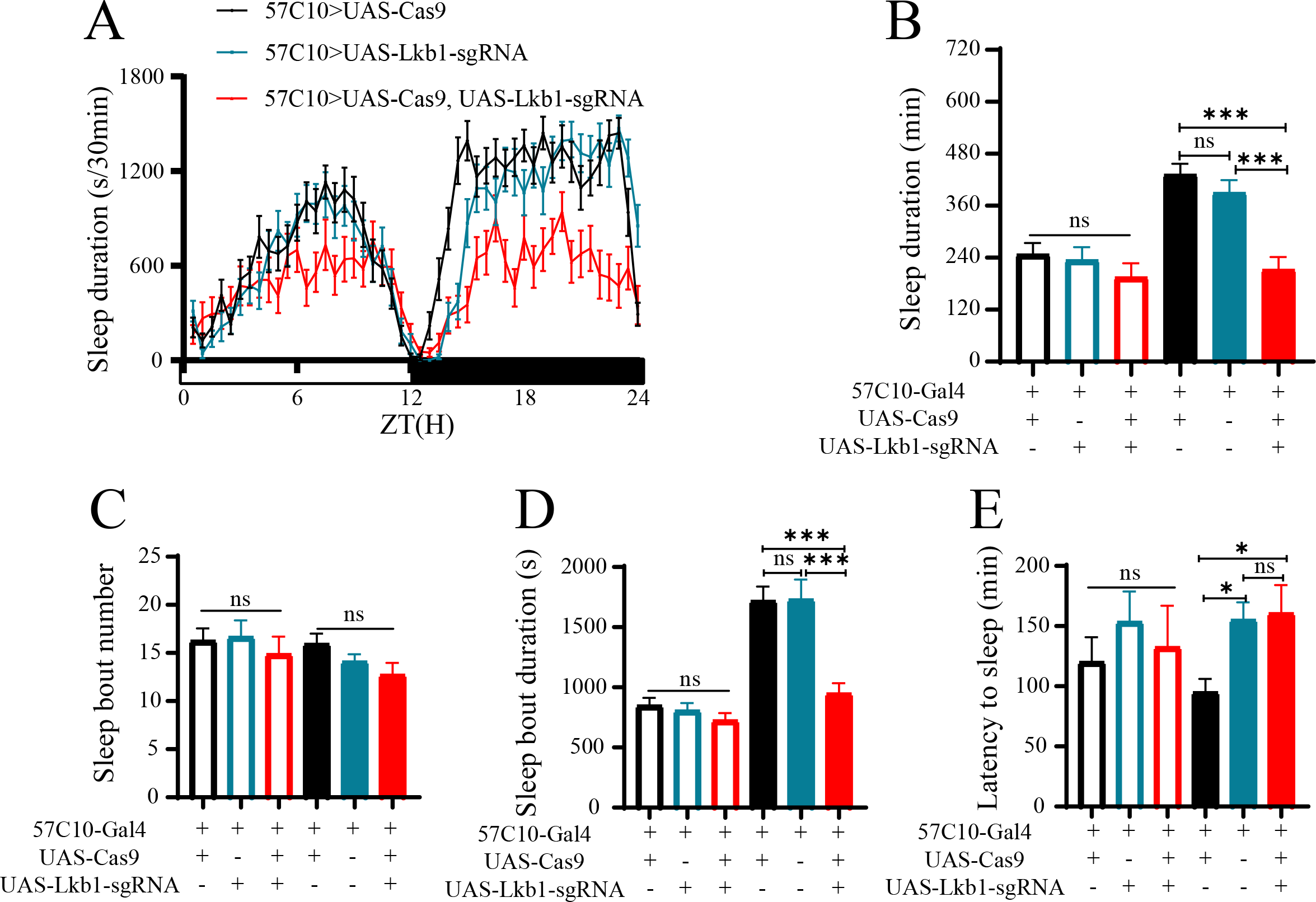
**Sleep Phenotypes of Mutants from Whose Neurons Lkb1 Was Targeted.** (A) Sleep profiles of UAS-Lkb1-sgRNA/57C10-Gal4;+/UAS-Cas9 (red, n=22), UAS-Lkb1-sgRNA/57C10-Gal4 (blue, n=24), and 57C10-Gal4/+;+/UAS-Cas9 (black, n=24) flies. (B-E) Statistical analysis of sleep duration, sleep bout number, sleep bout duration and latency to sleep in UAS-Lkb1-sgRNA/57C10-Gal4;+/UAS-Cas9 (red, n=22), UAS-Lkb1-sgRNA/57C10-Gal4 (blue, n=24) and 57C10-Gal4/+;+/UAS-Cas9 (black, n=24) flies. Open bars denote daytime, filled bars nighttime. (B) Sleep duration. Nighttime sleep duration of UAS-Lkb1-sgRNA/57C10-Gal4;+/UAS-Cas9 was significantly less than those of UAS-Lkb1-sgRNA/57C10-Gal4 and 57C10-Gal4/+;+/UAS-Cas9 flies. (C) Sleep bout number. Daytime and nighttime sleep bout numbers of UAS-Lkb1-sgRNA/57C10-Gal4;+/UAS-Cas9 was not significantly from those of UAS-Lkb1-sgRNA/57C10-Gal4 and 57C10-Gal4/+,+/UAS-Cas9 flies. (D) Sleep bout duration. Nighttime sleep bout duration of UAS-Lkb1-sgRNA/57C10-Gal4;+/UAS-Cas9 was significantly less than that of UAS-Lkb1-sgRNA/57C10-Gal4 and 57C10-Gal4/+,+/UAS-Cas9 flies. (E) Latency to sleep. Latency to sleep after light-off of UAS-Lkb1-sgRNA/57C10-Gal4;+/UAS-Cas9 was longer than that of 57C10-Gal4/+;+/UAS-Cas9 which was not significantly different from that of UAS-Lkb1-sgRNA/57C10-Gal4 flies. Ordinary One-way ANOVA was used. n.s. denotes p>0.05, *p<0.05, **p<0.01, ***p<0.001. Error bars represent SEM.

Daytime sleep duration, bout number, bout duration and latency were not significantly affected by neuronal gene targeting of Lkb1 (Figure 3B, 3C, 3D and 3E).

In all three series of experiments (Figures 1, 2 and 3), nighttime sleep phenotypes were more obvious than daytime sleep phenotypes. These results strongly indicate that Lkb1 expression in neurons are required physiologically for sleep, especially nighttime sleep.

### Sleep Phenotypes in Lkb1 Conditional Knockout Mice

To study the involvement of Lkb1 in sleep of mammalian animals, we obtained Lkb1^fl/fl^ mice in which the loxP sites flanked exons 3 to 6 of the Lkb1 gene (Nakada et al. 2010). To delete the Lkb1 gene from these mice, we injected adeno-associated viral (AAV) constructs expressing either the Cre recombinase together with the green fluorescent protein (GFP) or GFP alone to infect the mouse brain. Cre-GFP or GFP was under the control of a neuronal specific promoter hsyn (in AAV-PHP.B-hsyn-Cre-GFP or AAV-PHP.B-hsyn-GFP).

We analyzed the expression of LKB1 protein in mice (Figure 4A, 4B). Injection of Cre-GFP expressing virus into wt or Lkb1^fl/+^ mice did not reduce LKB1 protein expression in the brain. Neither did injection of only GFP expressing virus into Lkb1^fl/fl^ mice. This conclusion was reached by examination of either several mouse brains combined (Figure 4A), or individual mouse brains (Figure 4B).

**Figure 4.**
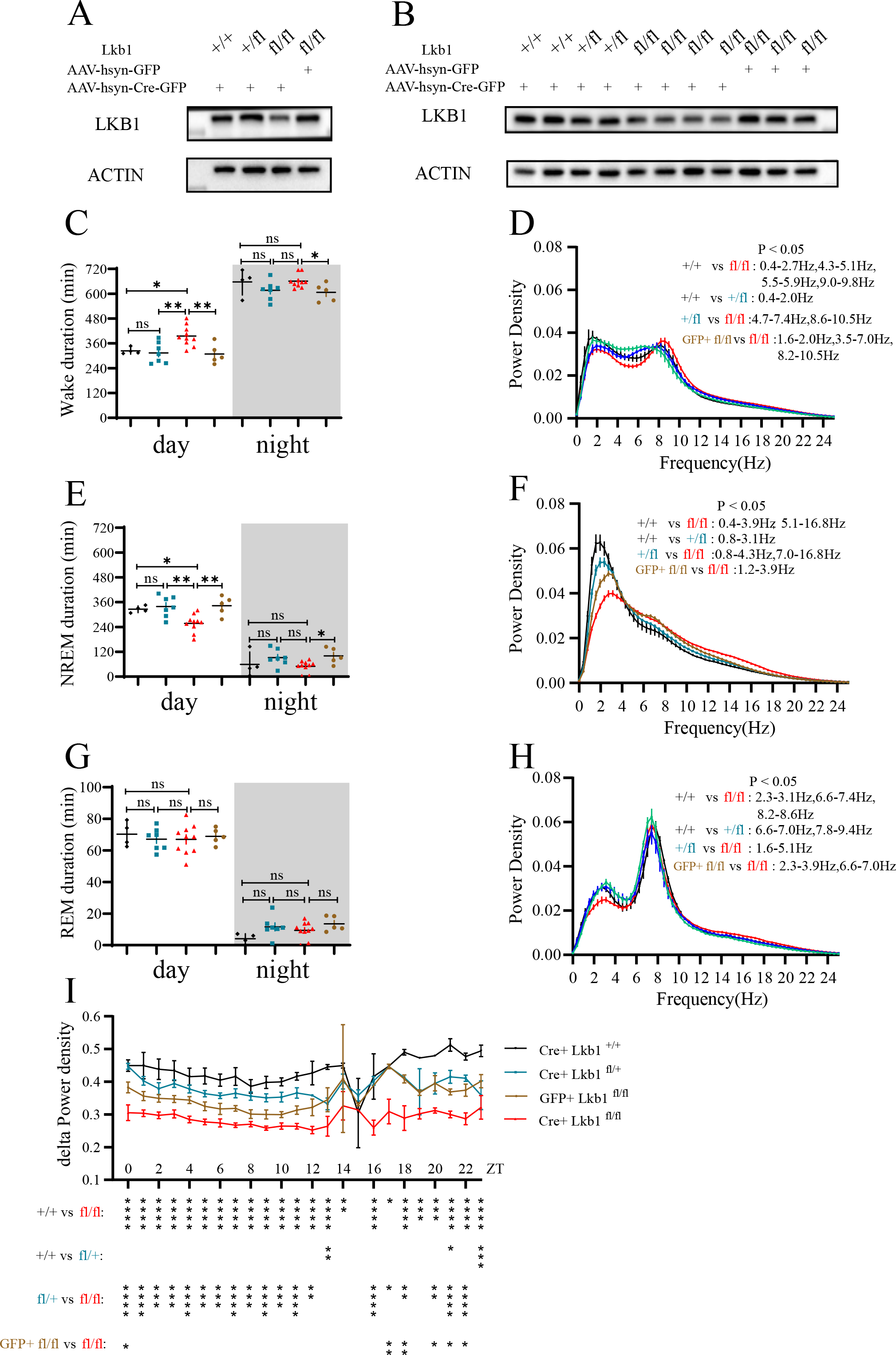
**Sleep Phenotypes of LKB1 Conditional Knockout Mice.** (A) Levels of LKB1 protein from Lkb1^fl/fl^ mice injected with AAV-hsyn-Cre-GFP virus (Cre^+^ Lkb1^fl/fl^, n=4), Lkb1^fl/fl^ mice injected with AAV-hsyn-GFP virus (GFP^+^ Lkb1^fl/fl^, n=3), Lkb1^fl/+^ mice injected with AAV-hsyn-Cre-GFP virus (Cre^+^ Lkb1^fl/+^, n=2) and Lkb1^+/+^ mice injected with AAV-hsyn-Cre-GFP virus (Cre^+^ Lkb1^+/+^, n=2). These mice were among those used for EEG recording and analysis. (B) Levels of LKB1 protein in individual mice (genotypes labelled: Cre^+^ Lkb1^fl/fl^, GFP^+^ Lkb1^fl/fl^, Cre^+^ Lkb1^fl/+^ and Cre^+^ Lkb1^+/+^. These mice were the same mice as those in (A) but presented individually. (C, E, G) Statistical analysis of wake duration, NREM duration and REM duration in Cre^+^ Lkb1^fl/fl^ (red, n=10), GFP^+^ Lkb1^fl/fl^ (yellow, n=5), Cre^+^ Lkb1^fl/+^ (blue, n=7) and Cre^+^ Lkb1^+/+^ (black, n=4) mice in a 12:12 LD cycle. White background denotes daytime, gray background nighttime. (C) Wake duration. Daytime wake duration of Cre^+^ Lkb1^fl/fl^ mice was higher than those of GFP^+^ Lkb1^fl/fl^, Cre^+^ Lkb1^fl/+^ or Cre^+^ Lkb1^+/+^ mice. Nighttime wake duration of Cre^+^ Lkb1^fl/fl^ mice was higher than that of GFP^+^ Lkb1^fl/fl^ mice. (E) NREM duration. Daytime NREM duration of Cre^+^ Lkb1^fl/fl^ mice was lower than those of GFP^+^ Lkb1^fl/fl^, Cre^+^ Lkb1^fl/+^ and Cre^+^ Lkb1^+/+^ mice. Nighttime NREM duration of Cre^+^ Lkb1^fl/fl^ mice was lower than that of GFP^+^ Lkb1^fl/fl^ mice. (G) REM duration. Daytime and nighttime REM durations of Cre^+^ Lkb1^fl/fl^ mice was not significantly different from those of GFP^+^ Lkb1^fl/fl^, Cre^+^ Lkb1^fl/+^ and Cre^+^ Lkb1^+/+^ mice. (D, F, H) EEG power spectrum of (D) WAKE, (F) NREM and (H) REM states in Cre^+^ Lkb1^fl/fl^ (red, n=8), GFP^+^ Lkb1^fl/fl^ (yellow, n=5), Cre^+^ Lkb1^fl/+^ (blue, n=6) and Cre^+^ Lkb1^+/+^ (black, n=4) mice. (I) NREM δ-power density of Cre^+^ Lkb1^fl/fl^ (red, n=8), GFP^+^ Lkb1^fl/fl^ (yellow, n=5), Cre^+^ Lkb1^fl/+^ (blue, n=6) and Cre^+^ Lkb1^+/+^ (black, n=4) mice. Ordinary One-way ANOVA was used in C, E, G for comparison of Cre^+^ Lkb1^fl/fl^, Cre^+^ Lkb1^fl/+^ and Cre^+^ Lkb1^+/+^ mice. Unpaired t test was used in C, E, G for comparison of Cre^+^ Lkb1^fl/fl^ and GFP^+^ Lkb1^fl/fl^ mice. Two-way ANOVA followed by Turkey’s multiple comparisons test was used in D, F, H, I. n.s. denotes p>0.05, *p<0.05, **p<0.01, ***p<0.001. Error bars represent SEM.

Functionally, only when the Cre-GFP expressing virus was injected into Lkb1^fl/fl^ mice, wake duration was significantly increased during daytime (Figure 4C, S4A), non-rapid eye movement (NREM) sleep duration was significantly decreased during daytime (Figure 4E, S4B). Controls (Cre-GFP injection into wt or Lkb1^fl/+^ mice, GFP injection into Lkb1^fl/fl^ mice) did not significantly changed any sleep phenotypes.

Rapid eye movement (REM) sleep duration was not significantly affected by Cre-GFP injection into Lkb1^fl/fl^ mice (Figure 4G, S4C).

Power density in the 1-4 Hz range (δ power density) of NREM is a commonly accepted indicator of sleep need (Borbely 1982; Borbely et al. 1981; Daan et al. 1984; Dijk et al. 1987; Franken et al. 2001; Tobler and Borbely 1986; Werth et al. 1996). We found that NREM δ power density was significantly reduced when the Cre-GFP expressing virus was injected into Lkb1^fl/fl^ mice (Figure 4F). Analysis over 24 h indicated that significant reduction was observed over most of the daily cycle (Figure 4I).

### Circadian Rhythm in Lkb1 Mutant Flies

The transcription factor differentiated embryo-chondrocyte 1 (DEC1) regulates circadian rhythm and can negatively regulate the transcription of Lkb1 and subsequently reduce AMPK activity (Sato et al. 2015).

We tested whether the circadian rhythm was affected in Lkb1 mutant flies. Lkb1 mutant flies were not different from wt flies in period length (Figure S5). Relative rhythmic power was increased in lkb1^T2/+^ and lkb1^T2/T2^ mutants than wt flies. (Figure S5).

## DISCUSSION

Our results indicate that LKB1 is required for sleep regulation: it plays an important and conserved role by promoting sleep in both flies and mice. LKB1 plays this role in neurons in both species because gene targeting of Lkb1 in neurons led to reduction of sleep. In mice, with the additional advantage of EEG analysis, we find that LKB1 regulates sleep need as indicated by reduced NREM δ power density in Lkb1 knockdown mutant mice.

Sleep is important for animals. Sleep regulation is accomplished through two processes: circadian and sleep homeostatic (Borbely 1982; Borbely et al. 2016). The circadian clock regulates the timing of sleep, and homeostatic process regulates the sleep drive. Molecular mechanisms of the circadian clock have been revealed through research in Drosophila and other organisms (Allada and Chung 2010; Mohawk et al. 2012; Nitabach and Taghert 2008). Although many sleep-related genes have been identified in sleep regulation (Allada et al. 2017; Cirelli 2009; Jan et al. 2020), our understanding of the mechanisms underlying sleep homeostatic regulation remains limited (Allada et al. 2017; Donlea et al. 2017).

Multiple regions in Drosophila and mouse brains have been implicated in sleep regulation. In flies, sleep is regulated by several regions including: the ILNv and DN1 clock neurons which are important for circadian control of sleep (Chung et al. 2009; Kunst et al. 2014; Parisky et al. 2008; Shang et al. 2013; Shang et al. 2008; Sheeba et al. 2008), and the mushroom bodies (MBs), the dorsal of fan-shaped body (dFB), the ellipsoid body (EB) the pars intercerebralis (PI) and glia (Chen et al. 2015; Crocker et al. 2010; Donlea et al. 2014; Donlea et al. 2011; Foltenyi et al. 2007; Guo et al. 2011; Joiner et al. 2006; Liu et al. 2012; Liu et al. 2016; Park et al. 2014; Pimentel et al. 2016; Seugnet et al. 2011; Ueno et al. 2012; Yi et al. 2013). In mammals, sleep is regulated by monoaminergic, cholinergic, glutamatergic, and GABAergic neurons that are distributed in multiple regions including the brain stem, the preoptic hypothalamus, the lateral hypothalamus and the basal forebrain (Saper and Fuller 2017; Scammell et al. 2017; Weber and Dan 2016). It will be interesting to investigate whether Lkb1 functions in all or a limited subset of neurons to regulate sleep.

Lkb1 as a master kinase can regulate the activities of ARKs by phosphorylating the site in the active T loop equivalent to AMPK-T172 (Lizcano et al. 2004). Because both AMPK and SIK3 are involved in sleep regulation, it will be interesting to investigate downstream kinases mediating the function of Lkb1 in sleep regulation. Is it SIK3, AMPK, or other members of the AMPK related kinases? The Ca^2+^/calmodulin-dependent protein kinase kinase-2 (CaMKK2, also known as CaMKKβ) could phosphorylate AMPKα−172 (Anderson et al. 2008; Hawley et al. 2005; Hurley et al. 2005; Woods et al. 2005), but CaMKK2 could not phosphorylate the equivalent sites in the other AMPK related kinases, including SIK3 (Fogarty et al. 2010). It will be interesting to investigate whether and how CaMKK2 regulates sleep.

In Drosophila, LKB1 functions through SIK3 which phosphorylates histone deacetylase 4 (HDAC4) to regulate lipid storage (Choi et al. 2015). It will be interesting to investigate whether HDAC4 is downstream of LKB1 in sleep regulation.

More importantly, an important question for further studies is whether Lkb1 regulation of sleep is related to its regulation of metabolism. Changes in energy homeostasis directly and reversibly influence the sleep/wake cycle (Collet et al. 2016). Some molecules involved in metabolism regulate sleep (Bjorness and Greene 2009; Gerstner et al. 2011; Grubbs et al. 2020; Nixon et al. 2015; Taheri et al. 2004; Thimgan et al. 2010). In Drosophila, starvation suppresses sleep without building up sleep drive (Thimgan et al. 2010). Lkb1 and its downstream components are involved in regulating metabolism, with examples such as LKB1-AMPK signaling in the liver regulating glucose homeostasis (Shaw et al. 2005), SIK3-HDAC4 regulating energy balance in Drosophila (Wang et al. 2011). Either LKB1 has two independent roles in sleep and metabolism or that its two roles are related.

## MATERIALS AND METHODS

### Fly Lines and Rearing Conditions

Flies were reared on standard corn meal at 25℃, 60% humidity and kept in 12 hours light/12 h dark (LD) conditions. 57C10-Gal4, nos-phiC31, hs-Cre on X were from the Bloomington Stock Center. vas-Cas9 was a gift from Dr. J. Ni (Tsinghua University, Beijing). UAS-Cas9 was constructed by Renbo Mao in our laboratory.

Flies used in behavioral assays were backcrossed into our isogenized Canton S background for 7 generations. All results of sleep analysis in this paper were obtained from female flies.

### Generation of KO, KI and Transgenic Lines

Total RNA was extracted from isoCS by TRIzol reagent (Invitrogen). Using the PrimeScript II 1st Strand cDNA Synthesis Kit (Takara), we reverse-transcribed the extracted mRNA into cDNA. The UAS-Lkb1 flies was constructed by inserting the coding sequence of CG9374 amplified from cDNA into the pACU2 plasmid (a gift from the Jan Lab at UCSF) (Han et al. 2011) before being inserted into the attP40 site. The UAS-Lkb1-sgRNA construct was designed by inserting the sgRNAs into pMt:sgRNA^3XEF^ vectors based on pACU2, with rice tRNA separating the different sgRNAs. CRISPR-Gold website was used to design 3 sgRNAs of Lkb1 (Figure S3) (Chu et al. 2016; Poe et al. 2019). The construct was inserted into the attP40 site.

KO and KI lines were generated as described previously (Deng et al. 2019). Knockout flies were generated with the CRISPR/Cas9 system. Two different sgRNAs were constructed with U6b-sgRNA plasmids. The 5’ homologous arm and the 3’ homologous arm of ∼2kb amplified from the wt fly genome were inserted into a pBSK plasmid for homologous recombination repair. The cassette of attP-3P3-RFP was introduced in the middle. sgRNA plasmids and the donor plasmids were injected into vas-Cas9 embryos to introduce attP-3P3-RFP into the genome at the region of interest and replaced it by homologous recombination. 3P3-RFP served as a marker to screen for the correct flies. Primers across the homologous arms were designed to verify the sequences by PCR and DNA sequencing. attP site was introduced into the genome with 3P3-RFP-LoxP. For KI files, the nos-phiC31 virgin females were first crossed with knock-out males and the pBSK plasmid inserted with attB-T2A-Gal4-miniwhite-LoxP cassette was injected into the female embryos. Miniwhite serves as a marker to screen for the correct flies, which could be excised by the Cre/LoxP system. Primers were designed to verify the sequence by PCR and DNA sequencing.

### Quantitative PCR

Total RNA was extracted from 30 flies of 5-7 days old by TRIzol reagent (Invitrogen). The genomic template was removed using DNase (Takara). cDNA was reverse-transcribed using Takara’s PrimeScript II 1st Strand cDNA synthesis kit (Takara). Quantitative PCR was carried out with TransStart Top Green qPCR SuperMix kit (TransGen) in the Bio-Rad PCR system (CFX96 Touch Deep Well). The sequences of primers used to detect Lkb1 and RP49(endogenous control) mRNA are:

Lkb1-F: 5’ -GCCGTCAAGATCCTGACTA-3’

Lkb1-R: 5’-CTCCGCTGGACCAGATG-3’

Rp49-F: 5’-CGACGCTTCAAGGGACAGTATC-3’

Rp49-R: 5’-TCCGACCAGGTTACAAGAACTCTC-3’

### Drosophila Sleep Analysis

Drosophila sleep analysis was performed as described previously (Dai et al. 2019; Qian et al. 2017). 5-7 days old flies were placed in a 65mm x 5mm clear glass tube with one end containing food and the other end plugging with cotton. All flies were recorded by video-cameras. Before sleep measurement, flies were entrained to an LD cycle at 25℃, 60% humidity for at least two days, and infrared LED light was used to ensure constant illumination when lights off. Immobility longer than 5 minutes was defined as one sleep event (Hendricks et al. 2000; Shaw et al. 2000). Information of fly location was tracked and sleep parameters were analyzed using Matlab (Mathworks), from which dead flies were removed. Sleep duration, sleep bout duration, sleep bout number and sleep latency for each LD were analyzed. Each experiment was repeated at least three times.

### Drosophila Circadian Analysis

Flies were reared and recorded in the same condition as sleep assay as described in papers from our lab (Dai et al. 2021; Qian et al. 2017), except that the condition was constant darkness. 6-8 days activity was measured and calculated in ActogramJ (Klarsfeld et al. 2003). Rhythmic strength, power and period were calculated by Chi-square method.

### Immunoblot Analysis

Mouse brains were quickly dissected and washed with phosphate buffer saline (PBS) on ice. Lysis buffer (20mM HEPES, 10mM KCl, 1.5mM MgCl_2_, 1mM EDTA, 1mM EGTA, 1mM DTT, freshly supplemented with a protease and phosphatase inhibitors cocktail) were used to homogenize brains by homogenizer (Wiggens, D-500 pro) at 4℃. Brain homogenates were centrifuged at 14,000 revolutions per minutes (rpm) for 15 minutes at 4℃. The supernatant was transferred to a new microtube and quantified with bicinchoninic acid assay (Thermo Fisher, 23225). The supernatant was analyzed by SDS-PAGE and proteins were transferred to a nitrocellulose membrane (GE Healthcare, #BA85). Membranes were incubated for 1 h in a blocking solution (Tris-buffered saline (TBS) containing 0.1% Tween-20, 5% milk). Primary antibodies were anti-LKB1 (cell signaling, #3047) and anti-actin (Santa Cruz, sc-8342).

### Retro-orbital Injection in Mice

Mice were reared at controlled temperature and humidity conditions with 12 h light/ 12 h dark cycle. Food and water were provided ad libitum. Lkb1^fl/fl^ mice were from the Jackson Laboratory (JAX #014143). They contained loxP sites flanking exons 3-6 of Lkb1 gene (Nakada et al. 2010). AAV-PHP.B-hsyn-CRE-GFP and AAV-PHP.B-hsyn-GFP virus were from Chinese Institute for Brain Research, Beijing. All results of sleep analysis in this paper were obtained from female mice.

0.2 ml/10 g Avertin was injected intraperitoneally into the mice for anesthetization. Rodent eyes were protruded by gentle downward pressure to the skin on the dorsal and ventral sides of the eye. The operator inserted the needle beveled downward into the retro-orbital sinus at the medial corner of the eye (Yardeni et al. 2011). The AAV-PHP.B virus was injected for whole brain infection (Chan et al. 2017).

### Mouse Sleep Analysis

Mouse sleep analysis was described in a previous article from our laboratory (Zhang et al. 2018). Eight-week-old mice were selected for retro-orbital injection. One week after viral injection, EEG and EMG electrode implantation procedures were performed. Mice were allowed to recover for more than 5 days individually and placed in a recording cage and tethered to an omni-directional arm (RWD Life Science Inc.) with connection cable for 2 days of habituation before recording. EEG and EMG data were recorded with custom software at a sampling frequency of 200 Hz for 2 consecutive days to analyze sleep/wake behavior under baseline conditions. The recording chamber was maintained at 12 h LD cycle and controlled temperature (24-25°C). EEG/EMG data were initially processed by Accusleep (Barger et al. 2019) before manual correction in SleepSign^TM^ to improve accuracy. WAKE was scored as high amplitude and variable EMG and fast and low amplitude EEG. NREM was scored as high amplitude δ (1-4 Hz) frequency EEG and low EMG tonus. REM was scored as a complete silent of EMG signals and low amplitude high frequency θ (6-9 Hz)-dominated EEG signal.

For power spectrum analysis, EEG was subjected to fast Fourier transform (FFT) analysis with a Hamming window method by SleepSign^TM^, yielding power spectra between 0-25 Hz with a 0.39Hz bin resolution. Epochs containing movement artifacts were marked, included in sleep duration analysis but excluded from the power spectra analysis. Power spectra for each vigilance state represents the mean power distribution of this state during a 24-h baseline recording. The δ-power density of NREMS per hour represents the average of δ-power density as a percentage of δ-band power (1-4 Hz) to total power (0-25 Hz) for each NREM epoch contained in an hour.

### Statistics

All statistical analyses were performed with Prism 7 (GraphPad Software). Differences in means between samples larger than two groups were analyzed using ordinary One-way ANOVA. Unpaired t test was used for two groups comparison. Power spectrum between different lines was compared by two-way ANOVA followed by Turkey’s multiple comparisons test. Ns denotes p>0.05, * denotes p<0.05, ** denotes p<0.01 and *** denotes p<0.001 for all statistical results in this paper.

## SUPPLEMENTAL INFORMATION

Supplemental information includes 5 figures.

## ACKNOWLEDGEMENTS

We are grateful to Ping-ping Yan, Lan Wang and Yong-hui Zhang for fly rearing, to Wei Yang and En-xin Zhou for Drosophila video tracing programs, to members of the Rao lab for discussion, to the Bloomington Drosophila Stock Center for flies, to the Jackson Laboratory for mice, to CIBR, Peking-Tsinghua Center for Life Sciences, IDG/McGovern Institute for Brain Research at Peking University and Changping Laboratory for support.

## Legends for Supplemental Figures

**Figure S1.**
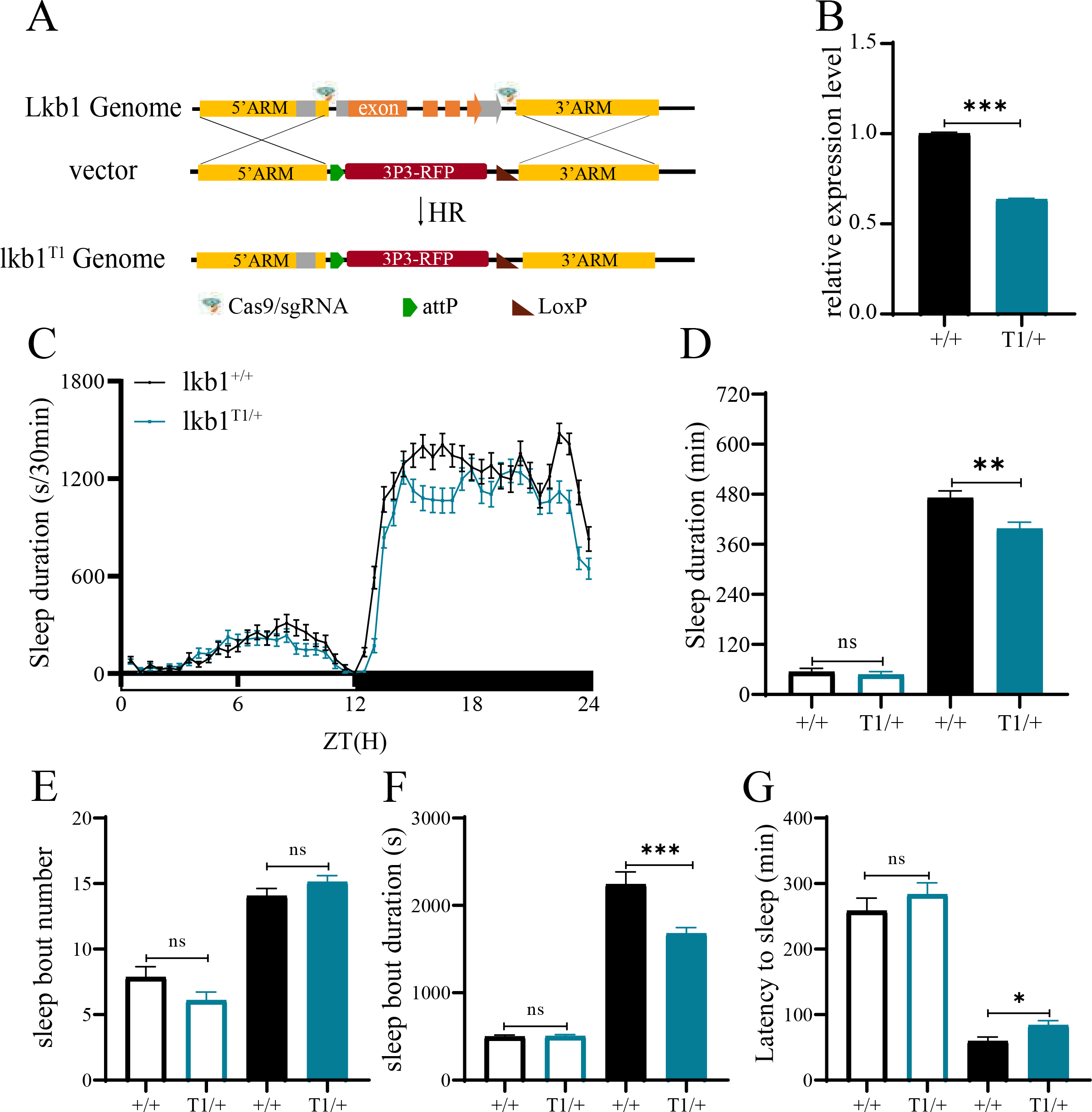
**Sleep Profiles of Lkb1 Knockout Mutants.** (A) A diagram representation of Lkb1 knock-out mutants lkb1^T1^. (B) Relative Lkb1 mRNA levels in lkb1^T1/+^ (blue) and wt (black) flies. lkb1^T1/T1^ was lethal. (C) Sleep profiles of lkb1^T1/+^ (blue, n=71) and wt (black, n=84) flies. (D-G) Statistical analysis of sleep duration, sleep bout number, sleep bout duration and latency to sleep in lkb1^T1/+^ (blue, n=71) and wt (black, n=84) flies. Open bars denote daytime, filled bars nighttime. (D) Sleep duration. Nighttime sleep duration of lkb1^T1/+^ was lower than that of wt flies. (E) Sleep bout number. Daytime and nighttime sleep bout numbers of lkb1^T1/+^ were not significantly from those of wt flies. (F) Sleep bout duration. Nighttime sleep bout duration of lkb1^T1/+^ was lower than that of wt flies. (G) Latency to sleep. Latency to sleep after light-off of lkb1^T1/+^ was longer than that of wt flies. Unpaired t test was used. n.s. denotes p>0.05, *p<0.05, **p<0.01, ***p<0.001. Error bars represent SEM

**Figure S2.**
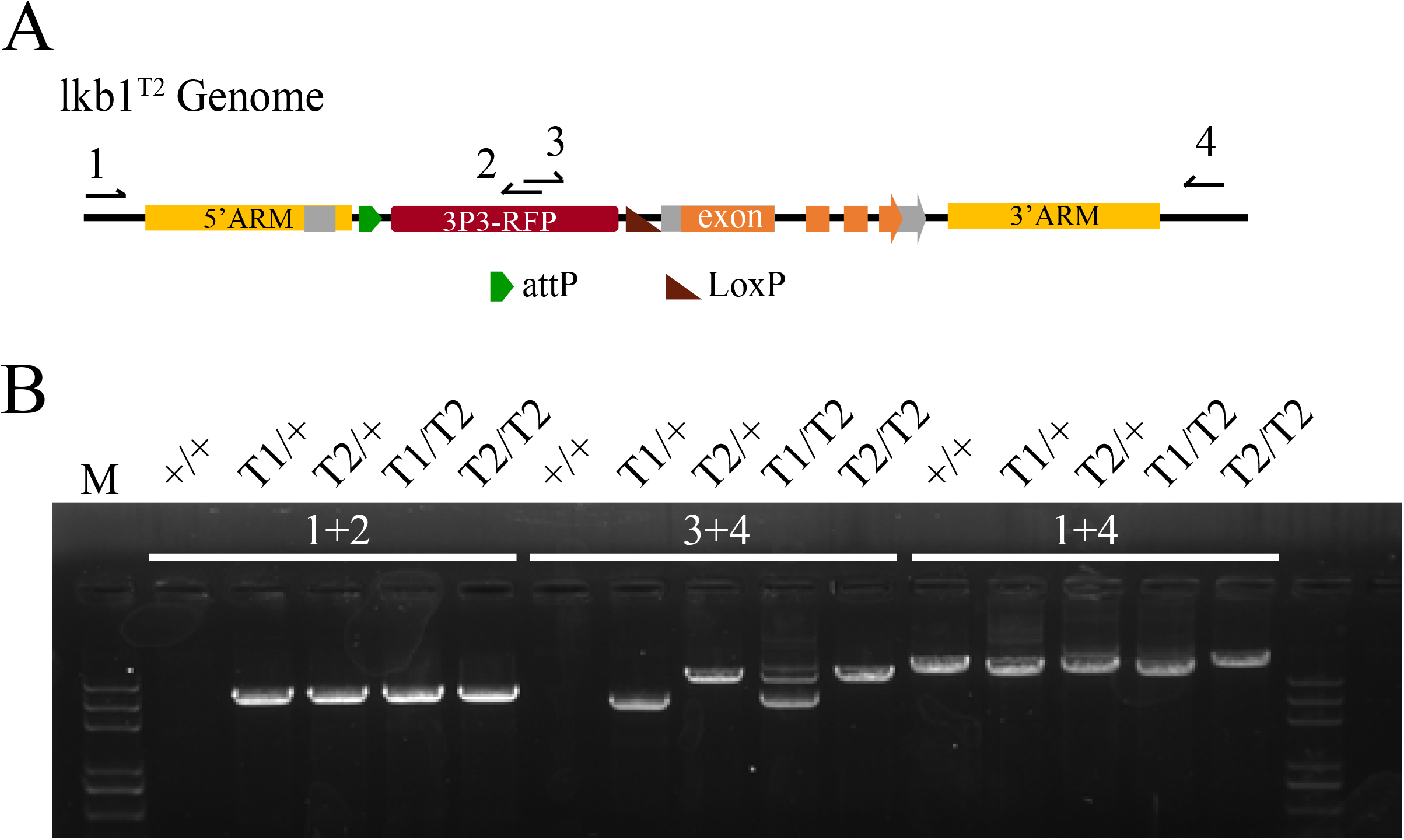
**Genotype Confirmation for lkb1^T2^.** (A) A diagram of lkb1^T2^ and the PCR primers used to detect inserted sequences. (B) Polymerase chain reaction (PCR) confirmation of the inserted sequences.

**Figure S3.**
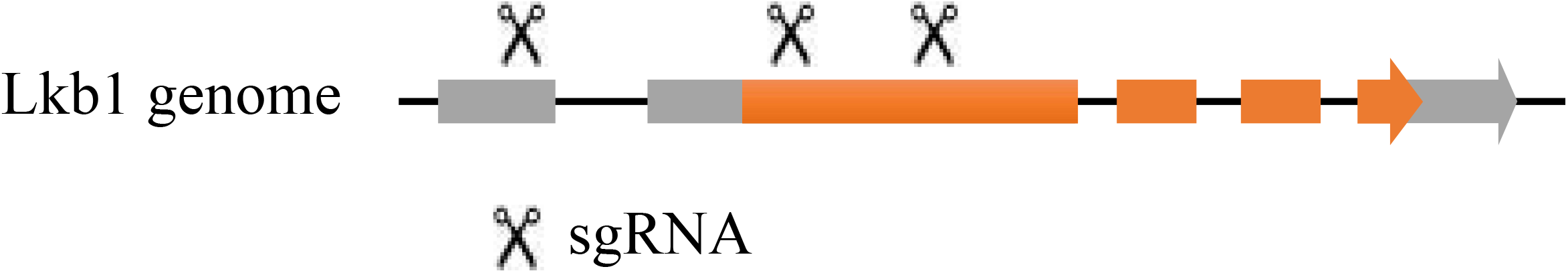
**Lkb1-sgRNA** A diagram of Lkb1 sgRNA. Three sgRNA were designed.

**Figure S4.**
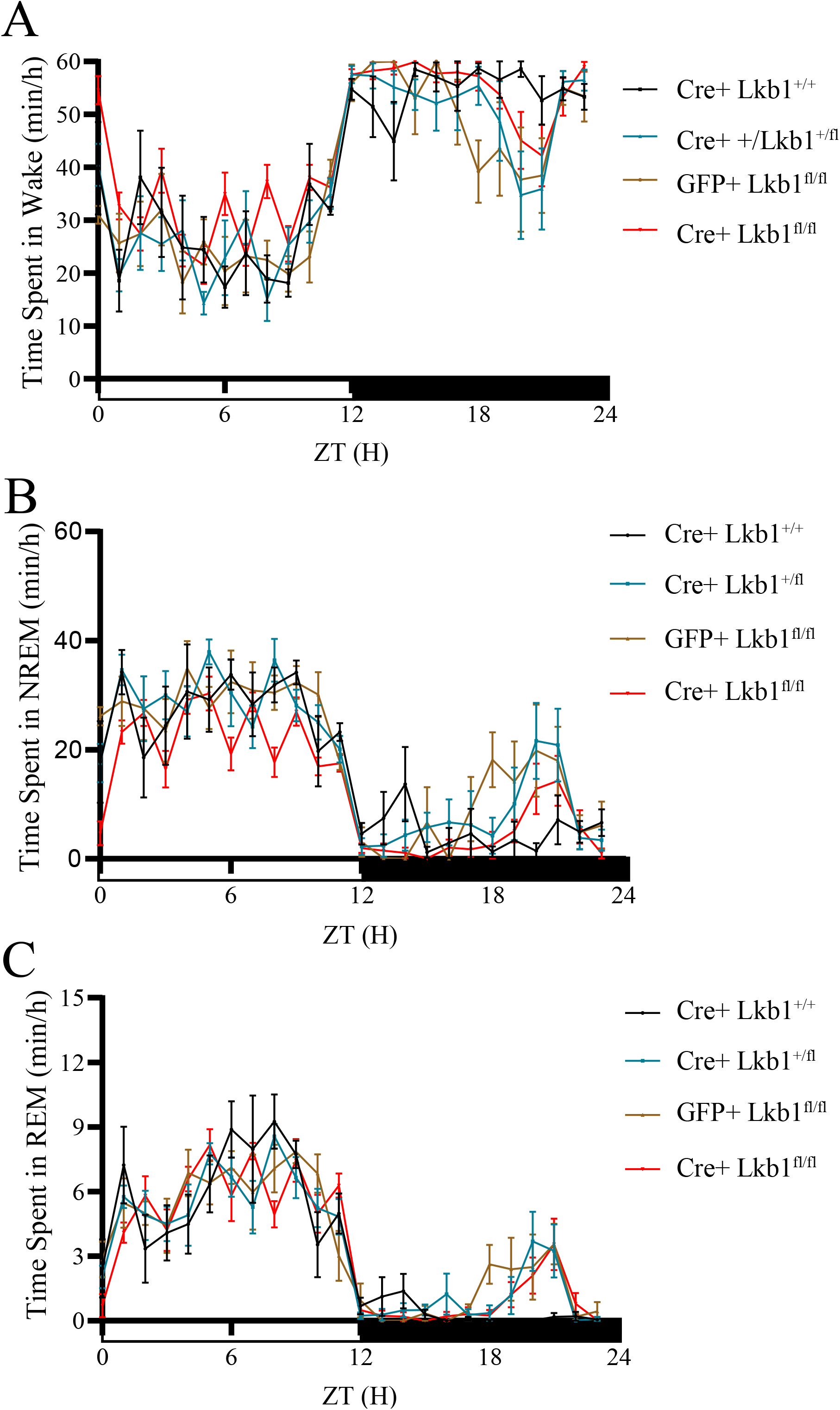
**Sleep Profiles of Lkb1-CKO in Mice.** (A) Time spent in wake of Cre^+^ Lkb1^fl/fl^ (red, n=10), GFP^+^ Lkb1^fl/fl^ (yellow, n=5), Cre^+^ Lkb1^fl/+^ (blue, n=7) and Cre^+^ Lkb1^+/+^ (black, n=4) mice in a 12 h light/12 h dark (LD) cycle. (B) Time spent in NREM of Cre^+^ Lkb1^fl/fl^ (red, n=10), GFP^+^ Lkb1^fl/fl^ (yellow, n=5), Cre^+^ Lkb1^fl/+^ (blue, n=7) and Cre^+^ Lkb1^+/+^ (black, n=4) mice in an LD cycle. (C) Time spent in REM of Cre^+^ Lkb1^fl/fl^ (red, n=10), GFP^+^ Lkb1^fl/fl^ (yellow, n=5), Cre^+^ Lkb1^fl/+^ (blue, n=7) and Cre^+^ Lkb1^+/+^ (black, n=4) mice in an LD cycle.

**Figure S5.**
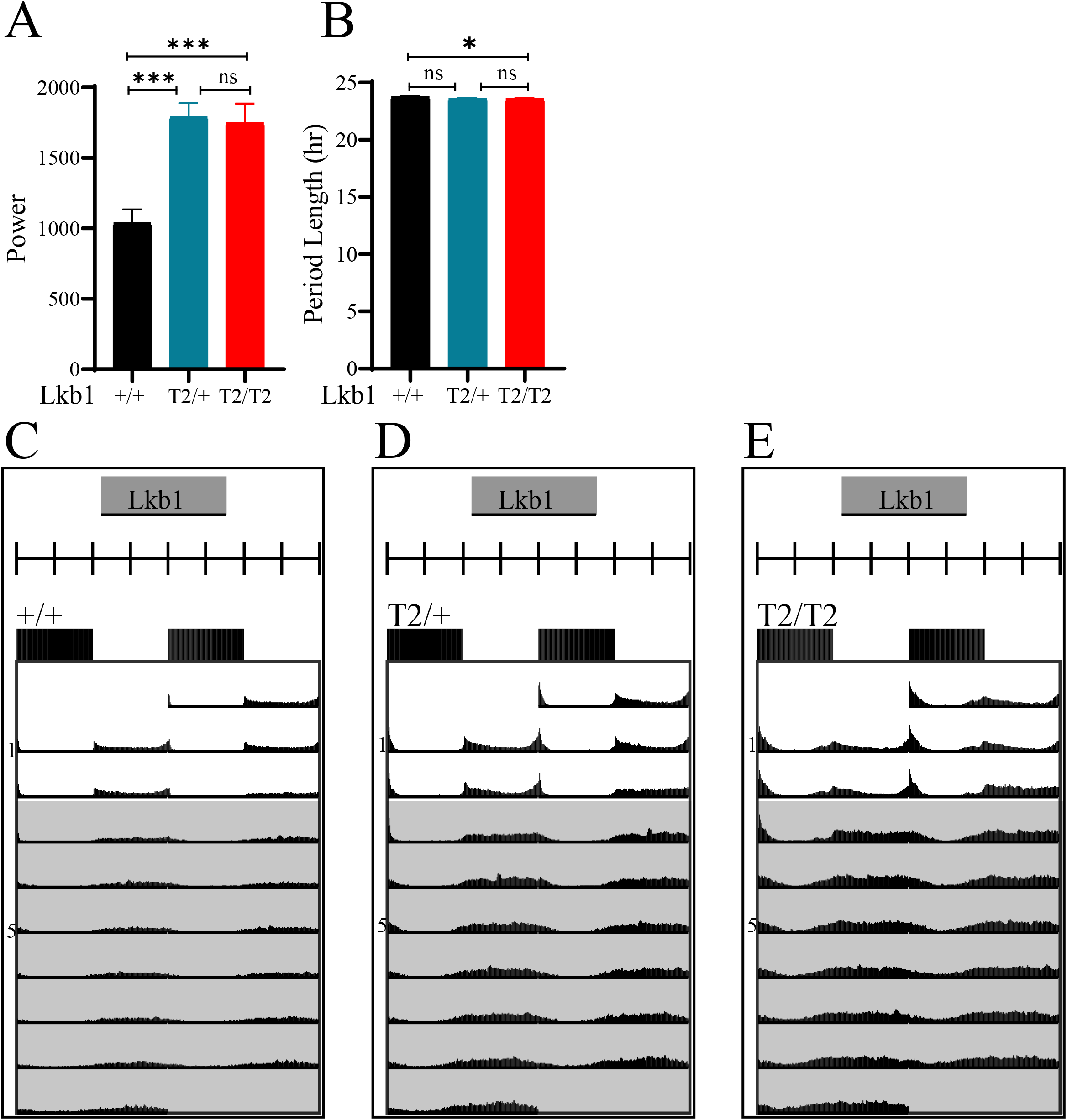
**Circadian Rhythm in Lkb1 Mutant Flies** (A) Relative rhythmic power in wt (black, n=42, 4/42 arrhythmic), lkb1^T2/+^ (blue, n=44, 0/44 arrhythmic) and lkb1^T2/T2^ (red, n=37, 0/37 arrhythmic) flies. The Power of lkb1^T2/+^ and lkb1^T2/T2^ were significantly higher than wt flies. (B) Period length in wt (black, n=42), lkb1^T2/+^ (blue, n=44) and lkb1^T2/T2^ (red, n=37) flies. The period of lkb1^T2/T2^ was not significantly different from those of wt and lkb1^T2/+^ flies. (C-E) Double-plotted actograms showing locomotor activity in wt (n=42), lkb1^T2/+^ (n=44) and lkb1^T2/T2^ (n=37) flies switching from 12hr LD to constant darkness. Open bars denote daytime, filled bars denote nighttime and grey background denotes DD. Ordinary One-way ANOVA was used. n.s. denotes p>0.05, *p<0.05, **p<0.01, ***p<0.001. Error bars represent SEM.

